# Searching for relief: *Drosophila melanogaster* navigation in a virtual bitter maze

**DOI:** 10.1101/804054

**Authors:** Nicola Meda, Giovanni Frighetto, Aram Megighian, Mauro Agostino Zordan

## Abstract

Animals use pain-relief learning to discern which actions can diminish or abolish noxious stimuli. If relief from pain is provided in a specific location, place learning is the mechanism used to pinpoint that location in space. Little is known about how physiological and non-directly damaging stimuli can alter visual-based searching behaviour in animals. Here we show how the optogenetically-induced activation of bitter-sensing neurons urges *Drosophila melanogaster* to seek relief from bitter taste stimulation and that this distressful, but ecologically relevant stimulus, innately wired to the threat of intoxication, is sufficient to elicit pain-relief-like behavioural responses. Specifically, freely walking flies inside an open circular arena are trained to seek relief from the unpleasant stimulation by searching for a safe area alternatively positioned in the proximity of a pair of identical, diametrically opposed, visual markers. Moreover, and perhaps more importantly, under this paradigm flies develop visual place learning manifested by their seeking relief in the zone associated with bitter relief during the last trial of training, even when exposed to constant bitter stimulation with no relief provided. An important implication is that this form of learning does not lead to operant conditioning generalization. We further propose that kinematic indexes, such as the spatially-specific reduction of locomotor velocity, may provide immediate evidence of relief-based place learning and spatial memory.

## Introduction

Animals are naturally prone to recognizing exogenous events or actions which can terminate or diminish noxious stimuli: this kind of learning ability is termed pain-relief learning (Navratilova and Porreca 2014; Gerber et al. 2014). The peculiarity of this type of learning lies in the fact that it depends upon the cessation of a negative stimulus rather than on the overt onset of a pleasant or unpleasant event as a consequence of an animal action (rewarding operant conditioning and punishing operant conditioning, respectively) (Quinn et al. 1974; Brembs 2000). While in the last two cases a rudimental receptivity towards external events may be sufficient to allow operant learning to take place, relief learning may depend on higher cognitive processes: an animal has to keep trace of the actions which do not lead to relief and employ other, more efficient behavioural strategies (Heisenberg 2015). Furthermore, the outcome seeked by the animal is solely the cessation of an internal state of distress, no other exogenous or environmental events are expected to occur. When relief from distress is provided in a specific place, the relieving action consists in being (and remaining) in the safe location (Baggett et al. 2018). Spatial localization based, for example, on visual (Ofstad et al. 2011) or idiothetic (Kim and Dickinson 2017) cues, is the learning mechanism that guarantees a faster return to the distress-relieving place in case of necessity (Ostrowski et al. 2015). The skills animals use to pinpoint relevant stimuli in the surrounding milieu is called place learning. If a specific location in space is associated to the cessation of noxious stimuli, such a location can be referred to, and be perceived by the animal, as a “safe zone” (Wustmann et al. 1996).

When the location of a fervently seeked safe zone is suddenly moved, the animal has to begin the searching behaviour anew in order to find the novel location. Most of research on this subject has relied on a visual-guided place-learning paradigm with heat (Ofstad et al. 2011), water (Morris 1984) or electrical shock (Yarali et al. 2008) acting as distress-inducing stimuli, and relief being provided by a safe location matched to a specific, unambiguous visual marker. Aside from directly-acting noxious events, little is known about how a specific physiological stimulus, with a negative valence, can alter search behaviours in animals. The way an animal alters its searching strategies in order to seek relief in an ambiguous visual environment has not, to our knowledge, been investigated yet.

Here we used optogenetic stimulation of bitter-sensing neurons in *Drosophila melanogaster* as a stimulus with a negative valence. The visual landmarks surrounding the animal in the testing arena were based on the stimuli used in Buridan’s paradigm (Götz 1980), with only one of the two opposing stripes marking the safe zone in each of a set of repeated trials (Figure 1A). In this type of environment only one of the two identical visual markers is matched to a zone in which no bitter stimulation occurs (i.e. the safe zone), the other is a decoy. With this virtual “bitter-taste maze” we were able to observe behavioural responses to a negative physiological stimulus and to its cessation; the alternate matching of only one of the two identical stripes to relief helped us to assess how the searching strategies of the flies changed in the presence of ambiguous visual markers. Our study evidences how bitter-taste can guide relief-based visual place learning in *Drosophila melanogaster*, and that this learning is associated with altered kinetics of locomotion when the animal enters the zone where relief is expected.

**Figure 1.**
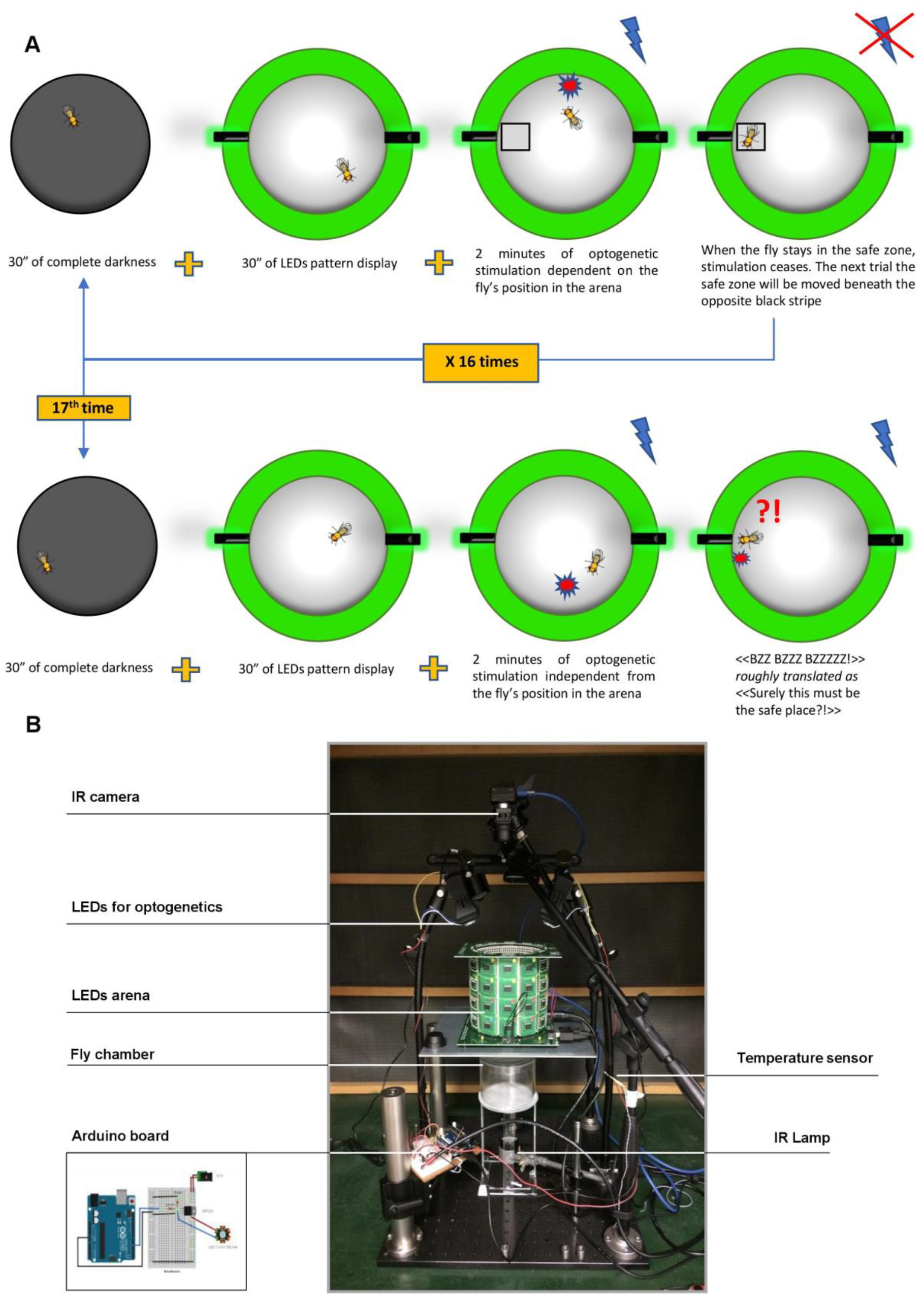
Experimental paradigm and apparatus used to investigate place learning in a virtual bitter maze. A single fruit fly at a time is tested. A) The first row of images depicts a single phase of a trial session. The fly is tracked in the arena for 3 minutes; during the first 30 seconds the fly is in complete darkness, for the following 30 seconds the fly is free to explore the surroundings with the visual pattern displayed; during the remaining 2 minutes optogenetic stimulation is delivered according to the fly’s position in the arena. This procedure is repeated 16 times in total. The end of the 16^th^ trial marks the end of the training session. After training (second row) the fly is probed for place memory in the same way as during a training trial, but without providing a safe zone. B) IR = infrared. For clarity we show the apparatus used in our experiments. With the exception of the optogenetic apparatus, the same tools were also used in a different set of experiments (Frighetto et al. 2019). Details about the set-up components are described in Materials and Methods section.

## Results

### Optogenetic stimulation of bitter-sensing neurons leads to relief-seeking behaviours in fruit flies

When exposed to electrical shock, heat, water or other distressing stimuli, animals do whatever they can to escape such a condition (Sneddon et al. 2014). While such stimuli can be directly damaging to an animal, little has been investigated on how bitter taste, a physiological and non-directly damaging stimulus, can alter the behaviour of freely-moving animals (König et al. 2014). In order to study whether bitter-taste could guide relief-based place learning, we used a *Gr66a-GAL4> UAS-XXL-ChR2* line expressing ChannelRhodopsin-2 (a blue-light sensitive rhodopsin) in most of the bitter-sensing neurons (from now on we will refer to this line as BitterStim). We first ran a training session of 48 minutes (16 trials each 3-minutes-long). Throughout this session, we tracked the behaviour of single flies online in a circular arena, surrounded by a grid of green LED panels (Figure 1B) displaying two identical and diametrically opposed vertical stripes (dark on a uniformly lit background), as employed in the classical Buridan paradigm. Flies were subject to constant optogenetic stimulation: from which relief was provided in a specific area of the arena, defined by a virtual “safe zone” (in the proximity of one of the two stripes). Whenever the tracked fly entered the safe zone, optogenetic stimulation was interrupted. At every trial, the safe zone was switched to the Buridan stripe which was not matched to relief in the immediately preceding trial. Using Generalised Linear Mixed Models (GLMM) to analyse the number of visits to the different zones of the arena and Linear Mixed-Effects Models (LME) to analyse the time spent in the different zones during each visit, we found that BitterStim and control flies differ in the number of visits to specific zones of the arena as well as in the time spent in those zones. In particular, BitterStim flies (N = 38) spent more time in the active safe zone than in any other arbitrarily defined zone having equal area and distance to the arena rim (Figure 2, Table 1, Supplemental_Movie_S1). The best GLMM explaining the number of visits to each zone takes into consideration the flies as belonging to different groups, whereas the best LME explaining the time spent in each zone is the one considering Group:Zone interaction as the explanatory variable (the best GLMM Bayes’ Factor – BF = 145.92 when compared to the null model; the best LME BF= 7.26*10^17^ when compared to the null model). We also observed that control flies (wild type Oregon-R strain (N = 8) and offspring of an Oregon-R x ChR2 (N = 10) cross) spent a slightly longer time in the active safe zone than in any other zone (Figure 2B), possibly due to the mildly distressing effect produced by the flashing blue-light (employed to produce the optogenetic stimulus).

**Table 1.**
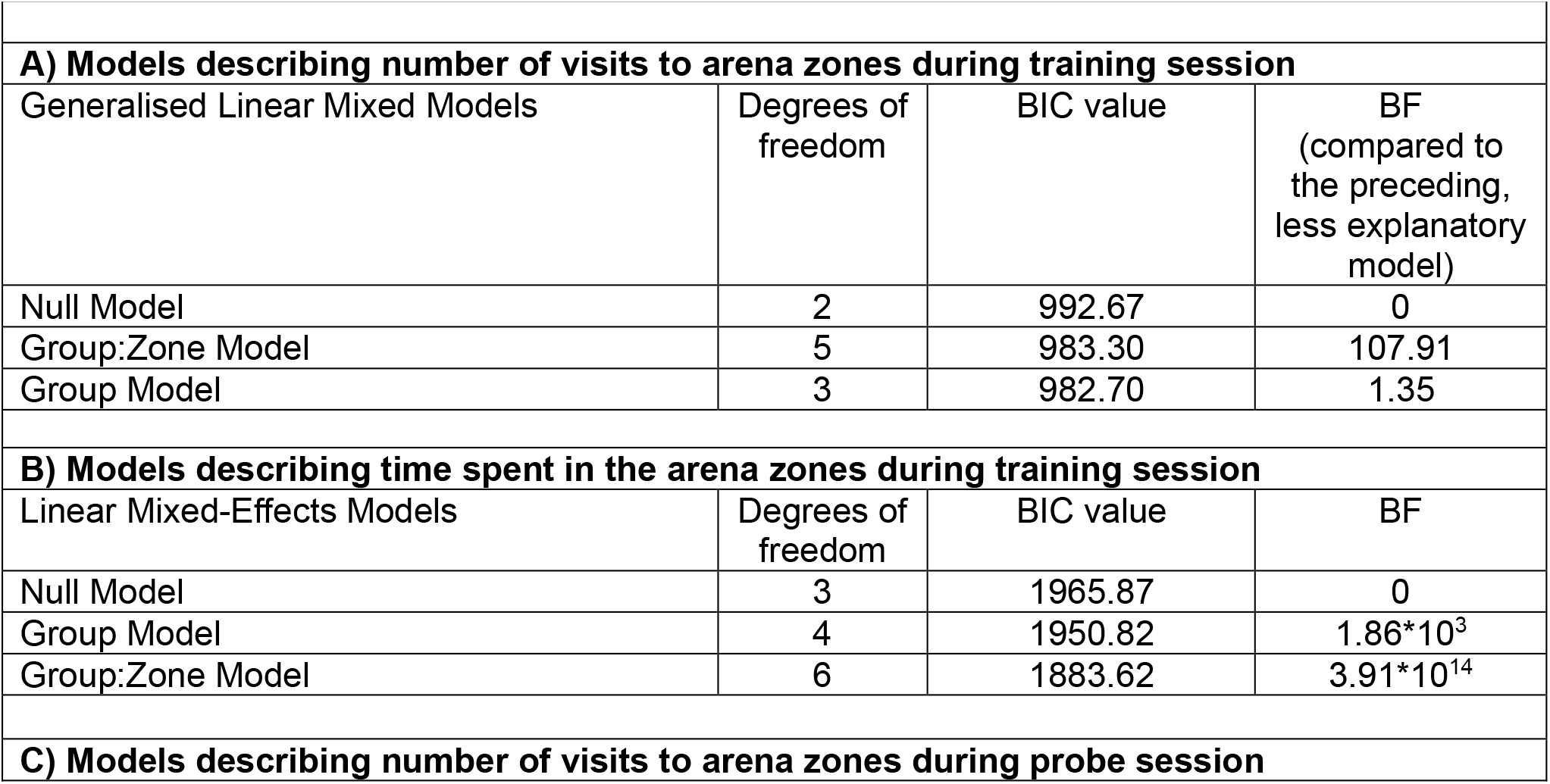
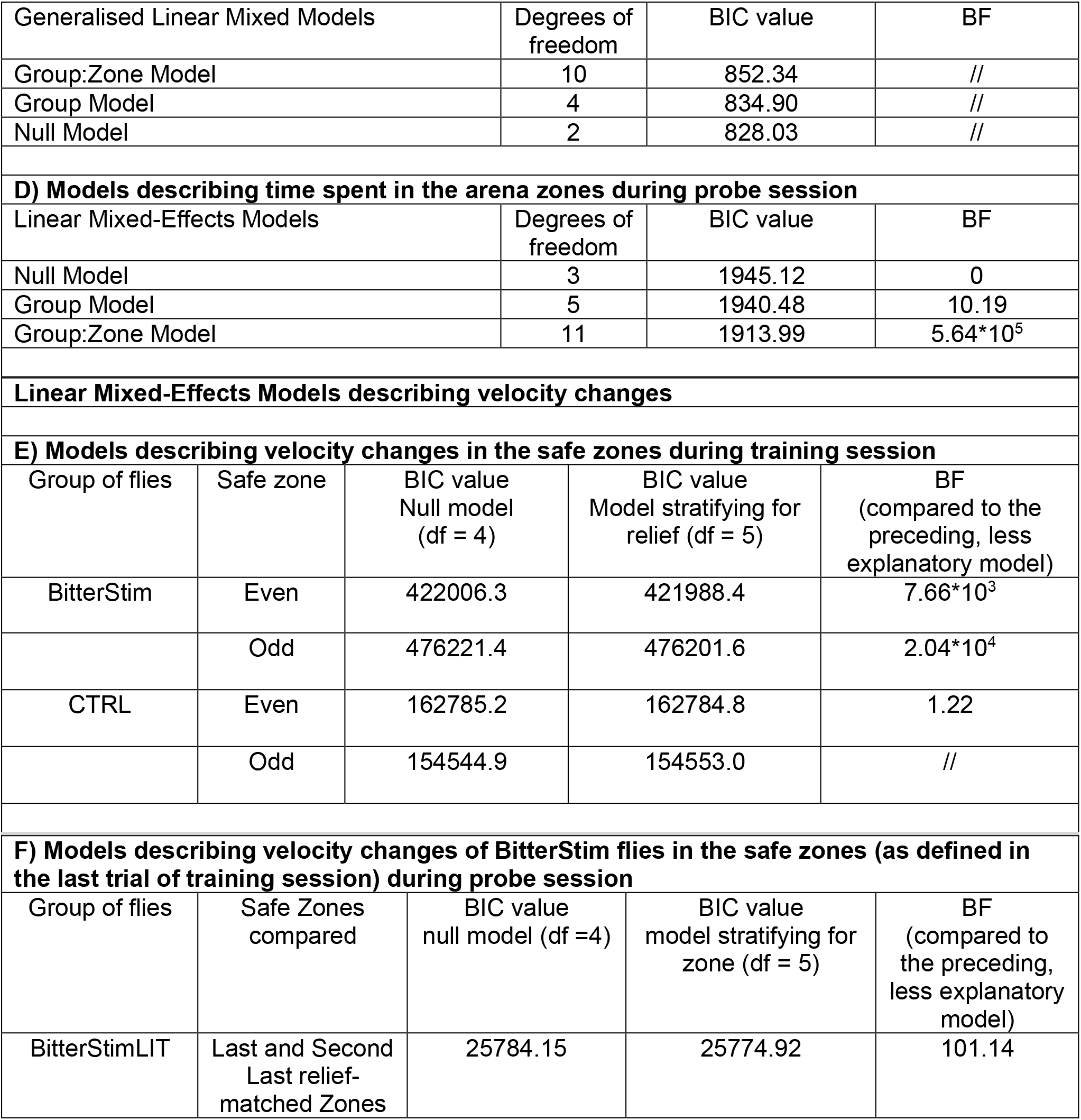
Linear Models describing the spatial distribution of flies and Linear Models describing locomotor velocity changes. BIC = Bayesian Information Criterion. BIC is an index of the goodness of fit of the model to the data, the lower the BIC the better the fit; BF = Bayes’ Factor; GLMM = Generalised Linear-Mixed Model; LME = Linear Mixed-Effects Model; df = degrees of freedom; Safe zone = relief-providing zone; Odd = southern safe zone, providing relief during odd numbered trials (framed by the sea-green square in Figure 2C); Even = northern safe zone, providing relief during even numbered trials (framed by the red square in Figure 2C); BitterStim = Bitter-stimulable flies; BitterStimLIT = Bitter-stimulable flies probed with LEDs displaying Buridan’s stripes lit; BitterStimDARK = Bitter-stimulable flies probed in complete darkness; CTRL = Control group. When the null model is the best model no BF is reported. A) Different GLMMs used to explain the number of visits to different zones in the arena during training session. In our case, the best GLMM for data explanation considers the individual flies as belonging to different groups, evidencing that there is a difference between bitter-stimulable flies and control flies; B) LMEs used to evidence a difference between group of flies in the time spent in different arena zones. Our analysis indicate that the best model for data explanation is the one considering the Group:Zone interaction as explanatory of the time spent in different zones. C) Different GLMMs used to evidence if flies inhomogeneous visits to different zones of the arena during the probe session can be explained by group and zone differences. Fruit fly visits to a zone of the arena during probe session are independent from the zone (whether it was a previously associated to relief or not) and the group to which the fly belongs. D) Different LMEs used to evidence that the time spent in specific zones of the arena during the probe session is dependent on Group:Zone interaction. E) Different Linear Mixed-Effects (LME) Models used to describe velocity changes during the training session. For the BitterStim group, velocity changes were better explained when considering whether a given zone, during specific trials, was an “active” safe zone (i.e. could provide bitter-relief). This means that the velocity curves, and therefore fly kinetics, depend on relief-perception. F) LMEs used to assess whether the velocity curves of BitterStimLIT flies are different between the last two relief-associated zones of the training session; BitterStimLIT flies manifested different velocities depending on the zone they were walking in.

**Figure 2.**
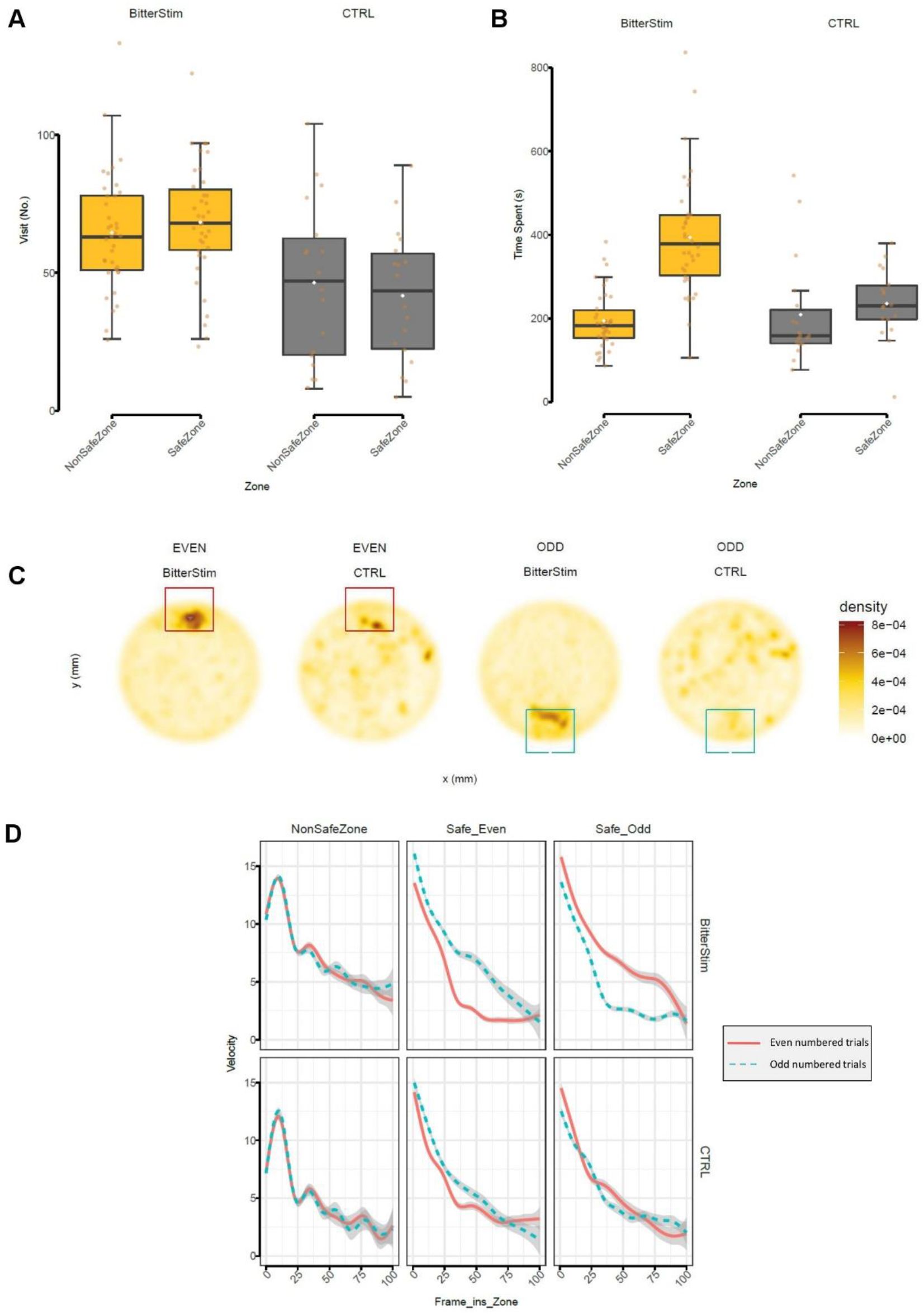
Bitter-stimulable flies explore the arena in search of relief during a training session and their locomotor velocity alteration is an index of relief perception. NonSafeZone = virtual zone designed for comparison; SafeZone = safe zone providing relief from optogenetic stimulation; BitterStim = Bitter-stimulable flies; CTRL = control strain; EVEN = when the safe zone is located on the “north” side of the arena (even numbered trials); ODD = when the safe zone is located on the “south” side (odd numbered trials); Safe_Odd = Safe zone active during odd numbered trials; Safe_Even = Safe zone active during even numbered trials; Frame_ins_Zone = progressive number of frames spent inside the zone; BIC = Bayesian Information Criterion; BF = Bayes’ Factor; df = degrees of freedom; GLMM = Generalised Linear Mixed Model; LME = Linear Mixed-Effects Model. A) Boxplots showing the number of visits to a specific zone, according to group. A visit is defined as the entrance in a computer-defined zone. The best GLMM in explaining the data, selected according to the lowest BIC, considers the two groups of flies as being different (in terms of their exploration): Group Model BF = 145.92, when compared to the null model. B) Boxplots displaying the time spent (in seconds) in specific zones of the arena by each group of flies. The time spent considers for how many frames a fly is located inside a computer-defined zone. The best model (LME) which explains the data considers the interaction between Group and Zone as explanatory of the time spent in each zone: BF = 7.26*10^17^, when compared to the null model. A) and B) Boxplots Boxes define first (Q1) and third (Q3) quartiles; bold horizontal black line is the median; white rhombus is the mean; brown circles are the values for single flies; whiskers define the lowest value still within 1.5 interquartile range [i.e., 1.5 × (Q3 − Q1)] of the lower quartile and the highest value still within the 1.5 interquartile range of the upper quartile. C) Density plots showing the zones most visited by flies (brownest areas). Plots are subdivided by group and by even/odd numbered trials. Density ranges from 8e-04 (brown) to 0 (white) for each group. During even numbered trials (the first two plots from the left), the safe zone is the one framed by the red square. During odd numbered trials (the last two plots from the left), the safe zone is the one framed by the sea-green square. BitterStim flies show a marked preference for the safe zone, compared to the more scattered distribution shown by control flies. D) For each group and zone two velocity curves are shown, as a function of the number of frames after the entrance into a specific zone (Frame_ins_Zone). The red continuous line represents the velocity (mm/s) curve during even numbered trials (i.e. when relief is provided in the “even” safe zone, framed by a red square in Figure 2C), the sea-green dashed curve represents the velocity during odd numbered trials. When the fly enters the active safe zone (e.g. the Safe_Even during even numbered trials), bitter-relief is provided. When such relief is experienced by the fly, velocity curve changes are detectable. BitterStim have significantly different velocity curves when optogenetic stimulation switches off and bitter-relief is provided, in both zones (Safe_Odd/South best LME – considering a relief effect – BF = 2.04*10^4^; Safe_Even/North best LME – considering a relief effect – BF = 7.66*10^3^). No significant difference in velocity kinetics is shown by control flies (CTRL) after cessation of the optogenetic stimulus (Safe_Odd best LME is the null model; Safe_Even best LME is the relief-effect model BF = 1.22, but this value is too low to be considered relevant according to (Raftery 1995)).

When a freely-behaving animal is exposed to a potentially harmful stimulus, it alters its behaviour in order to relieve distress and avoid any ensuing damage. Avoidance responses can be detected by considering the ensuing alteration in the kinetics of movement: for example, at the onset of an unexpected stimulus, independently from its valence, a startle response may be evoked (Bradley et al. 2018; Hoy et al.; Cho et al. 2004). Conversely, when relief from a noxious stimulus is associated with a particular spatial location, the animal might be expected to slow down and eventually stop in the safe zone; this may also be associated with a correlation between the degree of relief experienced and the magnitude of the decrease in velocity. Since, in our experimental paradigm, a specific safe zone provides relief on alternate trials, we compared the velocities of flies either experiencing or not experiencing relief in the same spatial location. A Linear Mixed-Effects Model shows that BitterStim flies present a more marked reduction in velocity when relief is provided, compared to when relief does not take place (the best LMEs (one for each relief-providing zone, i.e. north/even or south/odd) representing the BitterStim flies’ behaviour are those with relief as the explanatory variable; the northern safe zone BF= 7.66*10^3^ when compared to the null model; the southern safe zone BF = 2.04*10^4^ when compared to the null model). In the case of control flies, the best LME model for the southern/odd safe zone is the null model (i.e. with no explanatory variables), while we report a slight, and widely considered insignificant (Raftery 1995), difference for the northern/even zone (Table 1 and Figure 2D) with relief as the explanatory variable.

### Safe-place memory retention following training can be evidenced by localization and locomotor velocity reduction during a probe session

Following the training session, we wished to test whether or not flies had acquired a place preference. In order to do so, immediately after the training session, we ran a probe session which was designed to be similar to a single trial of training, but in this case no safe-zones were defined. We split the BitterStim group into two subgroups: one was probed in the arena in the presence of the Buridan stripes (BitterStimLIT, N = 21), the other was probed in complete darkness (BitterStimDARK, N = 20). BitterStimLIT flies, during optogenetic stimulation, are expected to search for the safe zones in the vicinity of the stripes more often than control flies (N = 18), as previously reported for similar visual learning paradigms (Ofstad et al. 2011; Collett et al. 1993). We employed the above described GLMMs and LMEs to test whether the frequency with which a group of flies entered specific zones of the arena, and the time spent in those zones, depended on bitter-stimulation. We found that a GLMM analysis for the number of visits to the arena zones, considering control, BitterStimLIT, and BitterStimDARK flies as belonging to different groups, did not highlight any significant differences, and the best model was a null model (Figure 3A and Table 1). No Group or Group:Zone interaction could explain the data, suggesting that, from the point of view of the number of visits to the arena zones, the three groups of flies were not statistically different. On the other hand, the best LME to describe the time spent within the arena zones by the flies was a model considering the interaction between Group and Zone. That is, BitterStimLIT, BitterStimDARK and control flies had different preferred zones in the arena (in terms of time spent within them) (Figure 3B and 3C, the best LME considers Group:Zone interaction as the explanatory variable for the time spent in different zones of the arena: BF= 5.75*10^6^ when compared to the null model).

**Figure 3.**
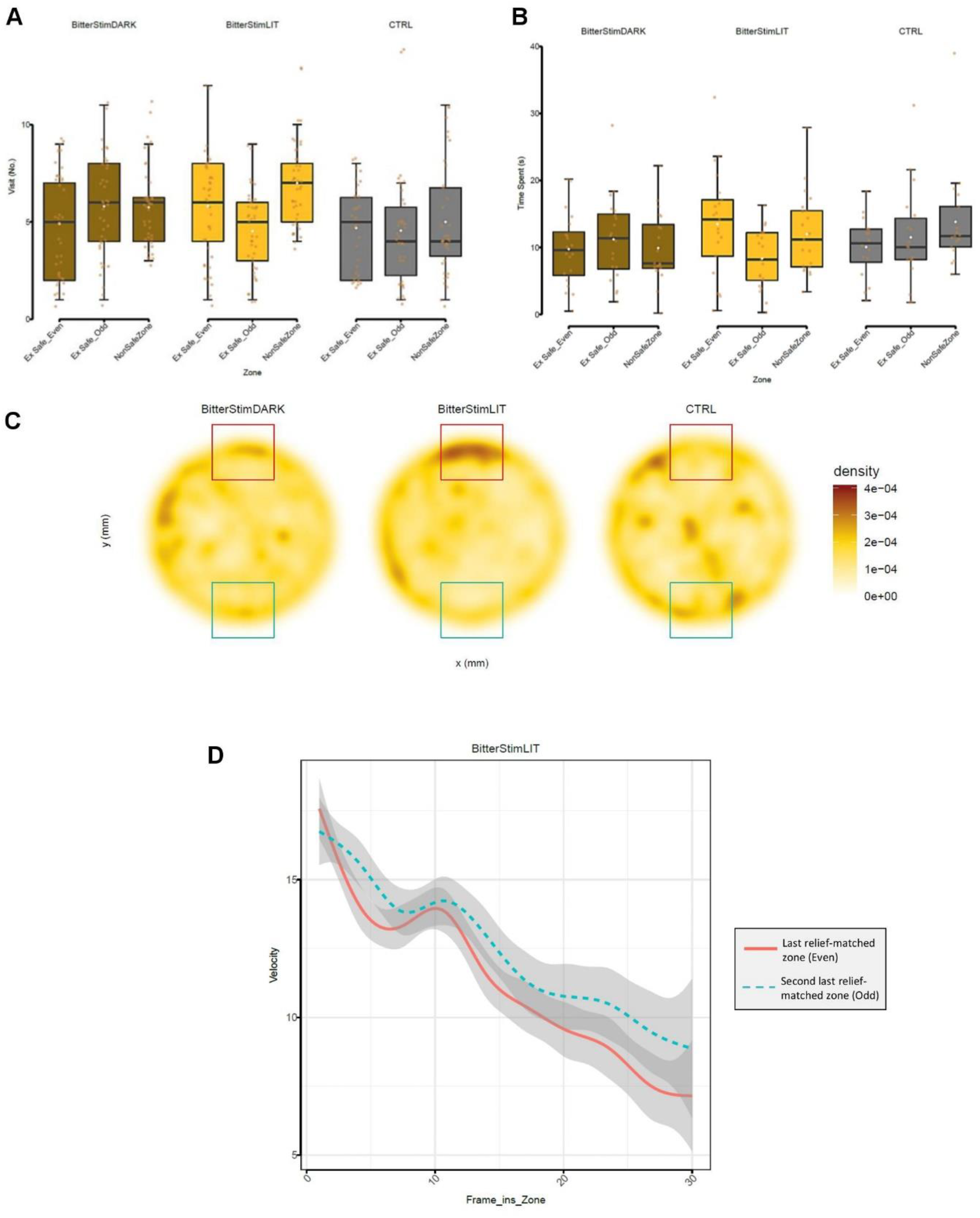
Bitter-stimulable flies probed with visual stimuli show a slight preference for one of the safe zones and their locomotor velocity alteration is concordant with this spatial observation. NonSafeZone = virtual zone designed for comparison; Ex Safe_Odd = southern safe zone, that provided relief during odd numbered trials of training (this is the second-last relief-matched zone); Ex Safe_Even = northern safe zone that provided relief during even numbered trials of training (this is the last relief-matched zone); BitterStimLIT = Bitter-stimulable flies probed with LEDs displaying Buridan stripes; BitterStimDARK = Bitter-stimulable flies probed in complete darkness; CTRL = control strain; Frame_ins_Zone = progressive number of frames spent inside the zone; BIC = Bayesian Information Criterion; BF = Bayes’ Factor; df = degrees of freedom; GLMM = Generalised Linear Mixed Model; LME = Linear Mixed-Effects Model. A) Boxplots showing the number of visits to a specific zone, according to group. A visit is defined as the entrance in a computer-defined zone. The best GLMM for data explanation is the null model, meaning that there are no differences in the number of visits in the different zones between groups. B) Boxplots displaying the time spent (in seconds) in specific zones of the arena by each group of flies. The time spent considers for how many frames a fly is located inside a computer-defined zone. Using a LME approach and best model selection according to the lowest BIC, we evidenced a difference in the time spent (s) in different arena zones that could be best described by a Group:Zone interaction model (BF= 5.75*10^6^ when compared to the null model). With particular reference to the BitterStimLIT group, we wanted to test if there were differences in the spatial distribution of flies between the Ex Safe_Even and Ex Safe_Odd zones. It turned out that the preference for Ex Safe_Even could still be evidenced by a LME model considering the two zones as not being homogeneously explored by the flies (BF= 186 when compared to the null model). This spatial feature can be seen also in the next panel. A) and B) Boxplots Boxes define first (Q1) and third (Q3) quartiles; bold horizontal black line is the median; white rhombus is the mean; brown circles are the values for single flies; whiskers define the lowest value still within 1.5 interquartile range [i.e., 1.5 × (Q3 − Q1)] of the lower quartile and the highest value still within the 1.5 interquartile range of the upper quartile. C) Density plots showing the most visited zones by flies (brownest areas). Density ranges from 4e-04 (brown) to 0 (white). BitterStimDARK flies (the first plot) show a diffuse searching behaviour, but no clear preference could be inferred. BitterStimLIT (the plot in the centre) seem to prefer the northern, Ex Safe_Even zone, even if no relief is provided. This effect could represent a manifestation of place preference. The CTRL group (plot on the right) shows no clear evidence of a place preference. D) Two velocity (mm/s) curves, for BitterStimLIT flies, are shown as a function of the time spent (in frames), one for each of the previous safe zones. The red continuous curve represents the velocity inside the last relief-matched (Ex Safe_Even) zone, the preferred zone for this group of flies; the sea-green dashed curve represents the velocity inside the second-last relief-matched zone. The best LME model describing the data takes into consideration that the two zones are different and evidences that BitterStimLIT flies had different velocity curves according to the zone considered (BF= 101.14 when compared to the null model).

Since during training both zones (i.e. the two diametrically opposed stripes) were alternately linked to bitter-relief, we expected BitterStimLIT flies to show approximately the same frequency of visits to each stripe during the post-training probe session. Surprisingly, this wasn’t the case: we found that BitterStimLIT flies manifested a safe zone preference (evidenced by the previous model and reported in Figure 3). In particular, these flies tended to prefer the stripe matched to relief in the last training trial. This did not occur for control flies.

From what we observed during training sessions, velocity could be used as an indirect index of place learning: that is, indirect evidence that an animal spends more time in a specific safe zone can be obtained by identifying bouts of decreased velocity, whenever relief can be achieved. As training progresses, an animal learns to expect specific consequences associated with its own actions or with external events (Heisenberg 2015). If a fly learns that stopping in the close vicinity of a visual landmark (i.e. Buridan stripe) can be relieving, this will lead to an alteration of the individual’s locomotor kinetics even before relief actually takes place. In our experiments, during probe sessions, no relief takes place but we reasoned that, following the 48 minutes of training, flies might form some kind of relief-expectation. To test this hypothesis we analysed the velocities of BitterStimLIT flies (the only group of flies that showed a marked preference for one of the two safe zones) when they were inside the previously (during training) safety-matched zones. We found that BitterStimLIT flies showed a significantly greater reduction in velocity when they were in the vicinity of the stripe which was associated to relief in the immediately preceding training trial than when they were close to the opposite stripe, suggesting that the two stripes yielded different mnemenic properties. (Figure 3D and Table 1. the best LME considers the two previous safe zones as being different also during the probe session: BF = 101.14).

### Visual markers are necessary for spatial preference

Animals can return to a known location by relying on multiple sources of information (Collett and Collett 2000; Wystrach et al. 2013; Gould 1986; David Morgan 2009). In our case, we wondered whether zone preference was based on idiothetic (i.e. due to path integration) or visual cues. To test this, a sub-group of bitter-stimulable fruit flies (BitterStimDARK) was probed in complete darkness (no Buridan stripes were displayed). Training scores (number of visits to the safe zones and time spent in the safe zones) for this sub-group were similar to those of the BitterStimLIT flies (Supplemental_Fig_S1). We then tested the location of the flies in the arena during the 30 seconds of complete darkness and during the 30 seconds before the onset of optogenetic stimulation. In order to avoid the confounding effect of flies already being present within a defined safe zone just before the onset of the optogenetic stimulation, we checked for this for all the test groups analysed and found only sporadic cases of flies being already present within a defined safe zone just before the onset of the optogenetic stimulation. In particular, no flies from the BitterStimLIT group were found to be in this condition (Supplemental_Fig_S2). When exposed to optogenetic stimulation during the probe session, BitterStimDARK flies neither visited the safe zones more often than the other arbitrarily defined zones having equal area and distance to the arena rim, nor, differently from BitterStimLIT flies, did they display a preference in terms of time spent in a particular safe zone. This feature supports the evidence that the place preference observed for BitterStimLIT flies is largely due to visual cues (Figure 3 and Table 1).

## Discussion

We used optogenetic stimulation of bitter-sensing neurons to study visual place learning in *Drosophila melanogaster*. Differently from previous paradigms for place learning (Ofstad et al. 2011; Yarali et al. 2008) our unconditioned stimulus was non-directly damaging and closely related to the ones used in path integration experiments (Corfas et al. 2019). Furthermore, two identical and diametrically opposed LED stripes were associated with relief from the bitter stimulus on alternate trials, thus adding an ambiguous visual cue-relief association. We showed that optogenetic activation of bitter-sensing neurons is distressing, and flies tend to seek relief from this stimulation alternatively between one of two identical stripes (Figure 2C). From an ethological perspective, each of the two stripes, which are identical, are linked alternatively either to relief or to the lack of relief. Thus, after the second trial of training, either stripe has been linked to bitter-taste relief or the lack thereof. In the latter case, if the negative bitter experience were to trigger an operant conditioning generalisation (i.e that in the vicinity of the black stripes bitter taste is not necessarily relieved), the flies’ bouts of relief searching would be directed away from the stripes. In nature, however, such a process would be detrimental to the animal: for instance, if a fly were to experience an instance of bitter-taste when visiting a red apple and this experience were generalised, the animal would cease to forage on red apples (or would do so with reluctance), thus excluding an important source of nourishment. In our experiments, even though the association between relief and zone is ambiguous (because of the inter-trial alternation) the recent memory of relief appears to override any generalisation. This is evident in the probe session: even if no relief can be achieved, BitterStimLIT flies remember the last location of the safe zone (Ex Safe_Even) and search in the vicinity of one of the stripes rather than navigating randomly in the whole arena. Bitter-relief is testified by two indirect behavioural signs of learning (Tomás Pereira and Burwell 2015): bitter-stimulable fruit flies spent more time close to a visually cued location yielding bitter-relieving properties than in other locations of the arena (Figure 2A, 2B and 2C); secondly, when bitter-stimulable flies experience bitter-relief, they slow down more quickly in order to stop in the safe zone (Figure 2D). This kinetic alteration is specific to bitter-relief: when the same spatial location does not provide relief, the velocity reduction is significantly less marked and does not necessarily lead to the fly stopping; furthermore, control flies do not alter their kinetics following the cessation of the optogenetic stimulation. Taken together, our results suggest that the preference which bitter-stimulable flies show for safe zones, during the training period, is attributable to bitter-relief. This behaviour is reminiscent of that elicited by heat or electrical shock punishment (Ofstad et al. 2011; Yarali et al. 2008). In line with these evidences, the threat of intoxication, innately wired to bitter taste, may be sufficient to elicit escape (in search of relief) responses (French et al. 2015) and alterations in locomotor velocity could be a marker of the relieving properties of a spatial location. Since the *Gr66a>Gal4* line we used for our experiments is characterised by expression of *Gal4* in almost all bitter-sensing neurons (Weiss et al. 2011; Freeman and Dahanukar 2015), more research on the subject is needed to assess the necessary and sufficient subset of bitter-sensing neurons which must be activated in order to elicit this complex response.

Throughout the training session (48 minutes), we trained flies to navigate between two identical visual landmarks (stripes) to experience relief from bitter stimulation. Thus, during the probe session, we expected flies to search for the safe zone approximately with the same frequency in the proximity of both Buridan stripes. Nonetheless, during the probe session, bitter-stimulable flies exposed to Buridan stripes (BitterStimLIT) manifested a searching behaviour preferentially centred on the stripe associated with relief during the last trial of the training session (Ex Safe_Even). This preference is characterised by two important features: BitterStimLIT flies spent more time in the Ex Safe_Even zone (Figure 3B and 3C) but also did not visit (i.e. enter) this zone significantly more than the other zones of the arena (Figure 3A and Table 1). This can be paraphrased as follows: even though BitterStimLIT flies did not frequently enter the computer-defined Ex Safe_Even zone, when they did they spent significantly more time searching for relief in that location than in the opposing Ex-Safe zone. This peculiar preference is also supported by the BitterStimLIT flies showing a significantly greater and steeper reduction in locomotor velocity when walking inside the Ex Safe_Even zone than in the opposing Ex-Safe zone (Figure 3D).

We interpret these results as additional evidence of retained place information and relief-learning, since none of the bitter-stimulable flies were inside that specific zone before the onset of optogenetic stimulation.

We wondered whether our group of flies developed this spatial preference using path integration strategies (Wang 2016; Heinze et al. 2018). To test this hypotheses, we probed a group of flies (BitterStimDARK) in complete darkness. This group had been trained exactly the same way as the BitterStimLIT group and both groups showed analogous training scores. If path integration played a role in the preference we observed, the effect of this preference should still be present in complete darkness. This was not the case: in complete darkness our BitterStimDARK group flies neither visited a previous safe zone more frequently nor spent more time in one of these zones. This suggests that visual cues are necessary to locate where the safe zones should be searched for.

From an ethological point of view, these results suggest that the threat of intoxication is sufficient to guide escape responses and relief-based place learning. Our results also suggest that the study of learning in freely-behaving animals may be facilitated by analysing, among other variables, variations in locomotor kinetics. More in general, extending our results to other paradigms and species, we suggest that sudden accelerations and increases in locomotor velocity may be considered as markers of startle-like responses, while decelerations/decreased velocities might be linked to relief. Opting for this approach could lead to the identification of learning processes without the need of running a dedicated probe session following the training.

## Materials and Methods

### Experimental Models

Optogenetically bitter-stimulable offspring resulted from crossing Blooming Stock Center (BSC) line 57670 (genotype: *w*[*]; *wg*[Sp-1]/*CyO*; P{w[+mC]=*Gr66a*-*GAL4*.1.8}1/*TM3*, *Sb*[1]), characterised by bitter-sensing neurons-restricted expression of Gal4 (Weiss et al. 2011), to a UAS-ChannelRhodopsin2 construct-bearing line (w[*]; *UAS*-*XXL*-*ChR2*/*CyO*:*GFP*), a kind gift of Professor Christian Wegener. Please note that this type of ChR2 does not require all-trans-retinal in order to be activated by 480nm light (Pauls et al. 2015; Dawydow et al. 2014).

For our behavioural experiments we opted to use only adult male flies. As control lines, we used wild-type Oregon-R (OR-R, obtained from BSC, line 2376) and offspring from crossing OR-R strain to the *UAS-ChR2* construct-bearing line. All OR-R x ChR2 offspring selected for experiments was *CyO*-negative.

Parental lines and progeny were reared on 12-15mL of standard corn meal medium in plastic vials (height 13cm, diameter 5cm). Parental lines (10 males and 5 virgin females for each cross – selected 1-2 days after birth) were kept in the same vial to mate for 4-5 days. Virgin females from ChR2 line were crossed to 57670 line males. In the OR-R x ChR2 crosses, OR-R were used as the source of virgin females. After 4-5 days of mating, the parental flies were moved to another clean vial to mate and lay further eggs, and this procedure was repeated 3-4 times.

Offspring birth was monitored every day (between 9.00 am and 11.00 am) and hatched adult individuals were selected under CO_2_ anaesthesia according to phenotypic markers. 5-6 adult male individuals of such selected offspring were put in the same vial. The vials were then put in a black TJENA box (Ikea, SW). 4 LEDBERG white LEDs (Ikea, SW) were glued to the cover of the box. To avoid blue-light diffusion (LeDue et al. 2016) from these LEDs, we applied a red theatre jelly (Neewer, USA, EAN: 0191073005013) in front of each one. Switching on and off of LEDs was controlled by a timer to ensure a 12:12 hour dark-light cycle with a precision of ±10 min. Vials were inspected on a daily basis to judge the state of health of the flies. If there were early signs of growing bacteria, flies were immediately transferred to another clean vial for the remaining days before testing. No antibiotics were used.

### Optogenetics apparatus

To study freely moving fruit flies without clipping their wings, we used a transparent resin arena covered with a glass cover (0.3mm thick), both having a diameter of 109mm. The arena presents a flat inner circular area of 5.5 cm diameter. In this area, the distance between the arena and the cover is 3.5mm; the arena then gently slopes upward towards the edge with an 11° degree slope, thus gradually reducing the distance between the cover and the base. This angle is necessary to avoid the presence of a fly-walkable “wall rim”. Arena and cover were designed with AutoCAD® 2015 and 3D printed using transparent resin by iMaterialise, BE.

The arena is placed on a raised platform; 15 cm underneath this platform, an infrared light-emitting LED lamp (LIU850A, Thorlabs Inc., USA) illuminates the arena surface. We placed 3 sheets of tracing paper between the arena and the lamp to diffuse and reduce the infrared light intensity. 36 cm above the arena an infrared-sensitive Chameleon 3 camera (CM3-U3-13S2C-CS-BD, FLIR, USA) was used to record fly movements. A cylinder composed of 48 panels, each consisting of a 8×8 grid of green (520nm) LEDs (JF-MR-FA0001 Rev B, IO Rodeo, USA), encircled the arena and was used to display visual stimuli. We controlled individual LEDs in order to display specific lighting patterns (i.e. Buridan’s paradigm stripes) by means of a MATLAB script, uploaded to the LED controller via SD Card. 27±1 cm above, and on opposing sides of the arena, 4-blue-emitting-leds (SP-08-B3, LuxeonStar, USA) per side were controlled by a programmable Genuino circuit board (UNO REV3, Arduino, IT), and delivered constantly 0.35mW/cm^2^ of blue light (measured with a Photodiode Power Sensor, model PM16-120, by Thorlabs, USA) optogenetic stimulation according to the position of the fly, which was being tracked live.

Each component of this set-up is placed inside a chamber which can be covered by a black cloth, thus isolating the whole setup from environmental light. The same hardware described here had also been used in (Frighetto et al. 2019) (Figure 1B).

To deliver time-specific optogenetic stimulation according to the position of the fly in the arena, a modified version of the Motion Based Multiple Object Tracking script by MathWorks® (https://it.mathworks.com/help/vision/examples/motion-based-multiple-object-tracking.html) was customized by us to meet our specific requirements. In particular, we used a live tracking system to record fly movements at 11 frames-per-second (during training sessions). The probe session was recorded at 10 frames-per-second.

Real-time fly coordinates were used by the script to drive the Genuino board’s output signals (and consequently the optogenetic stimulation) as follows: whenever the (x; y) coordinates of the centroid representing the fly were within a "safe zone" in the arena, the optogenetic stimulus was turned off; as soon as the fly left the safe zone, the stimulus was on.

### Behavioural experiments

1 to 5 hours after first daylight, adult male flies aged 8-10 days, were aspirated singularly from the vial and loaded into the arena. Temperature inside the box was assessed with a PTC-10 thermistor (NPI electronic, DE) and varied between 21-25°C degrees; Humidity inside the box varied between 35-45% (measured with a TH-50 probe by Hama, DE).

After loading the fly, the arena was placed on the raised platform, and the cylindrical LED display lowered around the arena. The lower row of cylinder LEDs was below the level of the arena in order to ensure the homogeneous diffusion of light. The whole apparatus was then covered with black cloth and the customised MATLAB Motion Based Multiple Object Tracking script executed.

Behavioural experiments consisted of two parts, a training session and a probe session. Each training session was made up of 16 trials. Each trial lasted 3 minutes: during the first 30 seconds the fly was free to explore the arena in complete darkness; for the next 30 seconds the fly could explore the arena with the green LED lights on. The LEDs were programmed to display two opposing stripes (according to Buridan’s paradigm, i.e. two diametrically opposed black stripes on an evenly lit background); during the last 2 minutes, the fly could be subjected to optogenetic stimulation according to its position in the arena, with the green LED visual patterns still present. Whenever the fly’s coordinates were inside a previously defined virtual safe area, no optogenetic stimulation took place; as soon as the fly left the safe area, the optogenetic stimulus (blue-LEDs) was automatically switched on, and bitter-sensing neurons stimulated. The safe zone was defined to be in close proximity to one of the two Buridan stripes. In each trial, the safe zone was shifted to be adjacent to the opposite Buridan stripe: during even numbered trials (1^st^, 3^rd^,…,15^th^) the safe zone was adjacent to the ‘northern end’ stripe; during odd numbered trials, the safe zone was adjacent to the ‘southern end’ stripe.

After approximately 48 minutes the probe session, in which no safe zones were defined, was started.

Each probe session lasted 4 minutes: the first minute was identical to a training session trial; while the last 3 minutes were characterised by constant optogenetic stimulation independently of the fruit fly’s position.

Once the probe session ended, the fly was aspirated from the arena and discarded; the arena was then cleaned with distilled water and dried with blotting paper. Flies were divided into two experimental groups: the controls (OR-R and OR-R x ChR2 offspring – respectively N = 8 and N = 10) and a “case” group called BitterStim (optogenetic-sensitive flies, 57670 x ChR2 offspring – N = 38). The case group was further subdivided into two subgroups for the probe session: the first subgroup of flies (BitterStimLIT, N=21) was probed in the presence of the Buridan visual stimuli; the second subgroup was probed in complete darkness (BitterStimDARK, N=20). Trained BitterStim flies were the same number of probed flies. Nonetheless, 3 training videos could not be tracked due to video acquisition issues.

### Statistical Analysis

Videos acquired with the infrared camera were first uncompressed with VirtualDub 1.10.4 software, then tracked with Ctrax 0.5.18 (Branson et al. 2009). Tracking errors were fixed using the MATLAB FixErrorsGUI Ctrax package, and per-frame statistics computed. Computed videos were transformed to .txt format with MATLAB 2018b. Using RStudio 3.5.3 we generated a dataframe for each experimental condition, and defined the localization of the safe zones in the arena. For subsequent comparisons and subsetting we also defined the localization of two extra zones, located at 90 degrees with respect to the safe zones, and equal to the safe zones in terms of area and distance to the arena rim. These two zones were merged and considered as one, unique “NonSafeZone”.

Training and probe sessions were analysed separately. For the training session, each session was split into two sub-sets: one composed of odd numbered trials (1^st^,3^rd^,..15^th^ trial) and the other of even numbered trials. This made the analysis of place preference easier since the position of the active safe zone depended on the trial number. Thus, the number of visits to/time spent in an “active” safe zone can at most last for half of the training session (the “Safe_Even” zone provides relief only during even numbered trials). To compare the preference between relief-providing zones and neutral zones of the arena, we applied the same time filter used for safe zones (i.e. filtering according to the evenness of trial number) also to the NonSafeZone (we remained conservative and selected the time filter that maximized the number of localization of flies in these neutral zones). Furthermore, to assess only relief or distress-related behaviours, we considered only the frames when optogenetic stimulation was present according to the flies’ position in the arena (i.e. the first 660 frames of each trial were discarded). To assess if bitter-stimulable flies entered the safe zones more often than the controls, we fitted the data with three different Generalised Linear Mixed Models (GLMMs) by using the R package *lme4* (Bates et al. 2019) and then we compared the models to select the one which best explained the data. Since "number of visits" is a counting variable, we used the Poisson family of distributions.

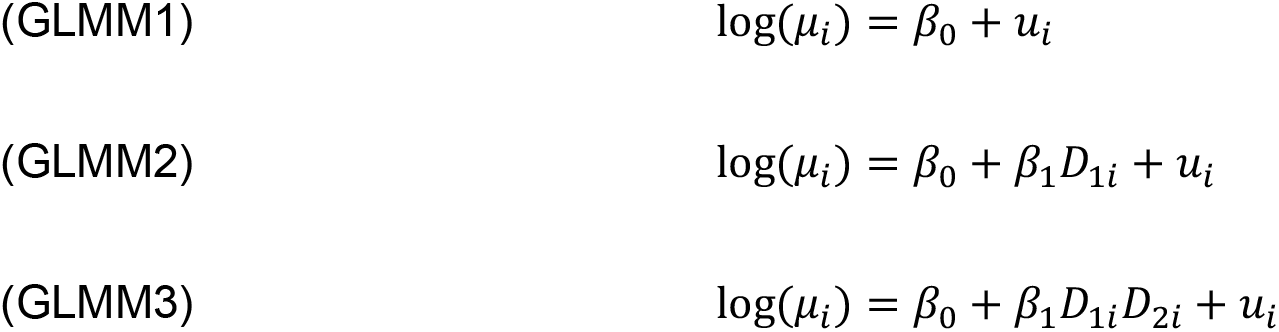

Where:

μ_*i*_ = incidence rate (number of visits)

*D*_*1i*_ = 1^st^ categorical predictor refers to group

*D*_*2i*_ = 2^nd^ categorical predictor refers to zone

*u*_*i*_ = random effects refer to the variation of the intercepts among files

To select the best fitting model we employed the Bayesian Information Criterion (BIC) (Schwarz 1978).

This approach was used to analyse the number of visits to arena zones of both training and probe sessions. To analyse place preferences during the probe session, we considered only the first two minutes of continuous optogenetic stimulation (the first minute and last minute of each probe session were cut, so as to allow the comparison of probe session data with those of the single trials during the training session). Since we considered the two safe zones as separate entities, we opted to filter out odd numbered frames from NonSafeZone detections, applying the same logic explained for the training session.

To assess if the time spent in different zones of the arena depended on the group of flies being tested and on the zones themselves, we fitted data with three different Linear Mixed-Effects Models (LMEs) by using the R package *nlme* (Pinheiro et al. 2018). BIC was the criterion employed to select the best model.

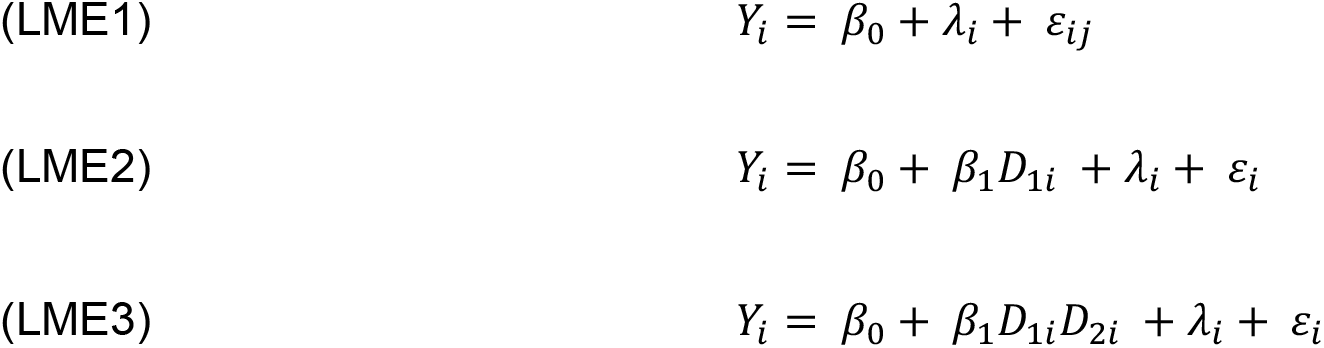

Where:

*Y*_*i*_ = *i*−_th_ linear model output (velocity)

*D*_*1i*_ = 1^st^ categorical predictor refers to group

*D*_*2i*_ = 2^nd^ categorical predictor refers to zone

*λ*_*i*_ = random effects refer to the variation of the intercepts among files

ϵ_*i*_ = error;

To test velocity changes within the safe zones, according to bitter-relief during training, we used a Linear Mixed-Effects Model. We considered a window of 100 frames (approximately 10 seconds) after the entrance of the fly into the zone: after ten seconds inside a zone, especially if relief is provided, a fly has probably stopped (so no further differences in velocity usually occur). We considered the progressive trials and flies as the nested random effects of the model (i.e. we let the intercept of the model vary for each fly during a specific trial). We computed two Linear Mixed-Effects Models: the first one was a “null” model, so velocity changes had only the progressive number of frames since the entrance in the zone as the explanatory variable. The second model considered velocity changes to vary as a function of relief being present or not and of the number of frames for which the fly was inside the zone.

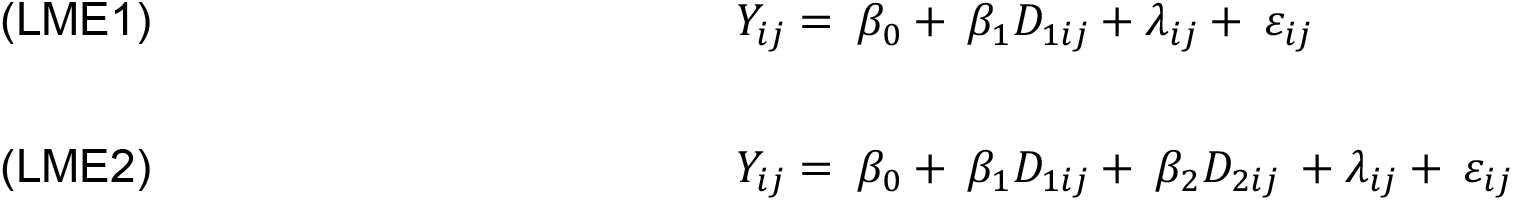

Where:

*Y*_*ij*_ = *i*−_th_ linear model output for the *j*−_th_ condition (velocity)

*D*_*1ij*_ = categorical predictor refers to progressive number of frames inside the zone

*D*_*2ij*_ = categorical predictor refers to relief (during training) or zone (during probe)

*λ*_*ij*_ = nested random effects (during training) or non-nested random effect (*λ*_*i*_)

ϵ_*ij*_ = error;

*D*_2*ij*_ in training session modelling is the relief condition for each fly (*i*) during a specific trial (*j*); *D*_2*ij*_ in probe session modelling is the zone condition for each fly (*i*) but the term *j* = 0 for all the variables in the equation (there is no nesting of random effects).

As for the GLMMs described previously, also for velocity changes, the BIC value guided model selection. An analogous approach was used to assess velocity changes during the probe session. In the latter case, we tested whether velocities were influenced by the zone (i.e. the safe zones) and progressive frame number of frames inside the zone, with random effects represented by each single experimental fly. In this case, since no relief is taking place anywhere in the arena, we hypothesised that BitterStimLIT flies would not stop inside any zone. So, no flies would remain within a zone, if they are not relieved from distress, for 10 seconds. Nonetheless, we opted to analyse the first 30 frames (approximately 3 seconds) as already done in similar paradigms applied to path integration (Corfas et al. 2019).

For graphics production, we used RStudio ggplot2 (Wickham 2009) and ggpubr (Kassambara 2018) packages. A step-by-step guide on analysis is provided as a comment in the RStudio script. Dataframes, MATLAB^®^ customised script and RStudio scripts are in Data_S1.

## Acknowledgments

We wish to thank Dr. Paola Cisotto for her unconditional technical support. We also thank Dr. Nicola Cellini for helpful suggestions.

